# An HIV-Tat inducible mouse model system of childhood HIV-associated nephropathy

**DOI:** 10.1101/2020.05.06.081851

**Authors:** Pingtao Tang, Jharna R. Das, Jinliang Li, Jing Yu, Patricio E. Ray

**Author notes:** Corresponding author: Patricio E Ray MD, Child Health Research Center, Room 2120, MR4 Building, 409 Lane Road, Charlottesville, VA. 22908.

## Abstract

**Background:** Modern antiretroviral therapies (ART) have decreased the prevalence of HIV-associated nephropathy (HIVAN). Nonetheless, we continue to see children and adolescents with HIVAN all over the world. Furthermore, once HIVAN is established in children, it is difficult to revert its long-term progression, and we need better animal models of childhood HIVAN to test new treatments.

**Objectives:** To define whether the HIV-1 trans-activator (Tat) gene precipitates HIVAN in young mice, and to develop an inducible mouse model of childhood HIVAN.

**Design/Methods:** An HIV-Tat gene cloned from a child with HIVAN was used to generate recombinant adenoviral vectors (rAd-Tat). rAd-Tat and *LacZ* control vectors (2 × 10^9^) were expressed in the kidney of newborn wild type and HIV-transgenic (Tg_26_) FVB/N mice without significant proteinuria (n = 5 - 8 per group). Mice were sacrificed 7 and 35 days later to assess their renal outcome, the expression of HIV-genes and growth factors, and markers of cell growth and differentiation by RT-qPCR, immunohistochemistry, and/or Western blots.

**Results:** HIV-Tat induced the expression of HIV-1 genes (*env*) and heparin binding growth factors in the kidney of HIV-Tg_26_ mice, and precipitated HIVAN in the first month of life. No significant renal changes were detected in wild type mice infected with rAd-Tat vectors, suggesting that HIV-Tat alone does not induce renal disease.

**Conclusion:** This new mouse model of childhood HIVAN highlights the critical role that HIV-Tat plays in the pathogenesis of HIVAN, and could be used to study the pathogenesis and treatment of HIVAN in children and adolescents.

**Summary statement:** We developed a new inducible mouse model system of childhood HIV-associated nephropathy, and demonstrated that HIV-Tat plays a critical role in this renal disease acting in synergy with other HIV-1 genes and heparin binding cytokines.

## Introduction

Modern combined antiretroviral therapies (cART) have improved the clinical outcome of children and adolescents living with HIV and decreased the prevalence of HIV-associated nephropathy (HIVAN) in a significant manner. However, physicians have had less success providing chronic cART to children and adolescents living with HIV, and we continue to see HIVAN in this group of people all over the world. Over 80% of the estimated 2.1 million HIV-infected children are living in the Sub Saharan Africa [1], and it is anticipated that approximately 10% of these children will develop HIVAN if they do not receive appropriate ART [1]. Furthermore, we have noticed that once the typical renal histological features of HIVAN are established in children, it is difficult to prevent its long-term progression to ESKD with current treatments available. In addition, previous reports in adults [2, 3] and children [4] suggest that HIVAN can occur in people with suppressed viral load. These studies suggest that inflammatory cytokines released by HIV-infected cells can play a role in the pathogenesis of HIVAN independently of the viral load. Taken together, all these findings underscore the importance of acquiring a better understanding of the pathogenesis and treatment of childhood HIVAN during the modern cART era.

Childhood HIVAN is a renal disease seen predominantly in Black children and adolescents who acquired HIV-1 through vertical transmission and do not receive appropriate antiretroviral therapy [5, 6]. From the clinical point of view it is characterized by persistent proteinuria, often in the nephrotic range, and in the late stages is associated with edema, reduced GFR, hypertension, and rapid progression to end stage kidney disease (ESKD) [1, 5, 6]. The renal histological lesions of childhood HIVAN reveal mesangial hyperplasia, focal segmental or collapsing glomerulosclerosis, and multicystic tubular dilatation leading to renal enlargement [1, 5, 6].

Several HIV-transgenic (HIV-Tg) animal models are available to study the pathogenesis and treatment of HIVAN [7–11]. However, these animals develop HIVAN at different time points, usually after they reach adulthood, and we lack a reliable mouse model system to study the pathogenesis of childhood HIVAN. Therefore, we carried out this study to determine whether the HIV-1 trans-activator (Tat) gene precipitates HIVAN in young mice, and define whether this approach could be used to generate an inducible mouse model system of childhood HIVAN. To accomplish this goal, we infected newborn wild type and heterozygous HIV-Tg_26_ mice with recombinant adenoviral vectors (rAd) carrying the coding sequence of the HIV-Transactivator gene (HIV-Tat) and assessed the renal outcome of these mice during the first month of life.

## Results

### Expression of HIV-Tat derived from a child with HIVAN in the kidney of newborn mice

The protein sequence of the HIV-Tat derived from a child with HIVAN (Tat-HIVAN) was aligned with Tat protein sequences derived from the lymphotropic virus HIV-1 IIIB and the monocyte-tropic HIV-1 virus ADA (NIH AIDS Research and Reference Reagent Program). As shown in Figure 1A, Tat-HIVAN contains the basic domain that is essential for HIV-1 activation, but is missing the RGD motif that interacts with cytokines and integrin receptors [12, 13]. Using an adenovirus gene transferring technique developed in our laboratory [14], we were able to express HIV-Tat in the kidney and liver of wild type and HIV-Tg_26_ mice (Figure 1B). As expected, by Western blots, higher Tat protein expression levels where detected in the liver compared to kidneys [14] (Figure 1B). HIV-Tg_26_ newborn mice infected with rAd-Tat showed higher Tat mRNA expression levels when compared to transgenic mice injected with rAd-*Lac-Z* vectors (Figure 1B). The Tat mRNA detected in HIV-Tg_26_ mice injected with rAd-*Lac-Z* vectors was transcribed from the HIV pro-viral DNA d1443 transgenic construct.

**Figure 1.**
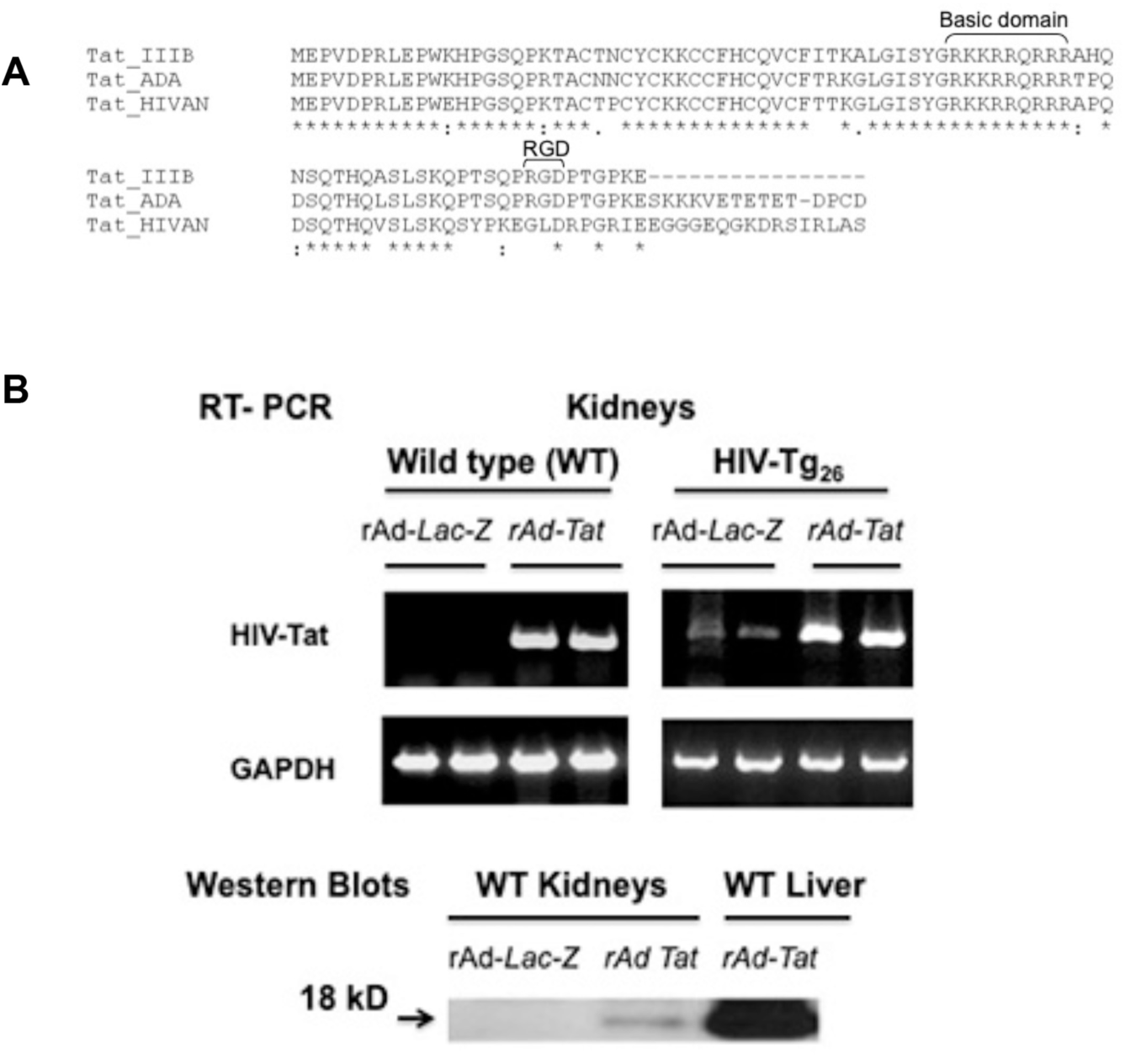
HIV-Tat gene expression in the kidney of newborn wild type (WT) and HIV-Tg_26_ mice infected with adenoviral vectors carrying Lac-Z or HIV-Tat coding sequences. **A.** The protein sequence of the HIV-Tat gene derived from a child with HIVAN is aligned with HIV-Tat derived from the lymphotropic HIV-IIIB virus or the monocyte-tropic HIV-1 virus ADA, using the Clustal Omega multiple sequence alignment program (http://www.ebi.ac.uk/Tools/msa/clustalo/). The basic domain and RGD motifs are indicated in brackets. The Tat sequencing procedure was repeated three times to rule out the possibility of a sequencing error. **B.** Newborn HIV-Tg_26_ mice were infected with recombinant adenoviral (rAd) vectors carrying either *Lac-Z (*rAd-*LacZ)* or HIV-Tat (rAd-Tat) vectors. Seven days later, all mice were sacrificed, and the kidneys were harvested and processed for the RT-PCR studies using specific Tat primers as described in detail in the methods section. The upper panel shows representative RT-PCR results corresponding to HIV-Tat mRNA expression in the kidney of young wild type (WT) HIV-Tg_26_ mice infected either with rAd-*LacZ* or rAd-Tat vectors. HIV-Tg_26_Tg mice infected with rAd-*LacZ* vectors showed Tat mRNA levels transcribed from the HIV-1 proviral DNA construct d1443 used to make the HIV-Tg_26_ mice. The lower panel shows representative Western blots Tat results in kidney and liver sections derived from wild type mice infected with rAd *Lac-Z* or rAd-Tat vectors.

### Tat-induced expression of HIV-genes, Fibroblast Growth Factor-2 (FGF-2), and Vascular Endothelial Growth Factor (VEGF-A) in young HIV-Tg_26_ mice

Seven days old HIV-Tg_26_ mice injected with rAd-Tat vectors showed higher renal expression levels of HIV-*envelope (env)* mRNA (~ 12 fold) by RT-qPCR (Figure 2A). The renal expression levels of HIV-*env* mRNA continued to be elevated (~ 5 folds) in 35 days old HIV-Tg_26_ injected with rAd-Tat vectors, when compared to those injected with rAd-*Lac-Z* vectors (Figure 2A). In addition, the mRNA expression levels of two heparin-binding cytokines (FGF-2 and VEGF-A) which are involved in the pathogenesis of HIVAN in HIV-Tg_26_ mice and children living with HIV [15–17], were elevated in the kidney of 7 days old HIV-Tg_26_ mice (Figure 2B-C). In contrast, lower expression levels of HIV-*env*, FGF-2, and VEGF-A were noted at 35 days. The later findings are consistent with the immune-mediated clearance of the Tat adenoviral vectors.

**Figure 2.**
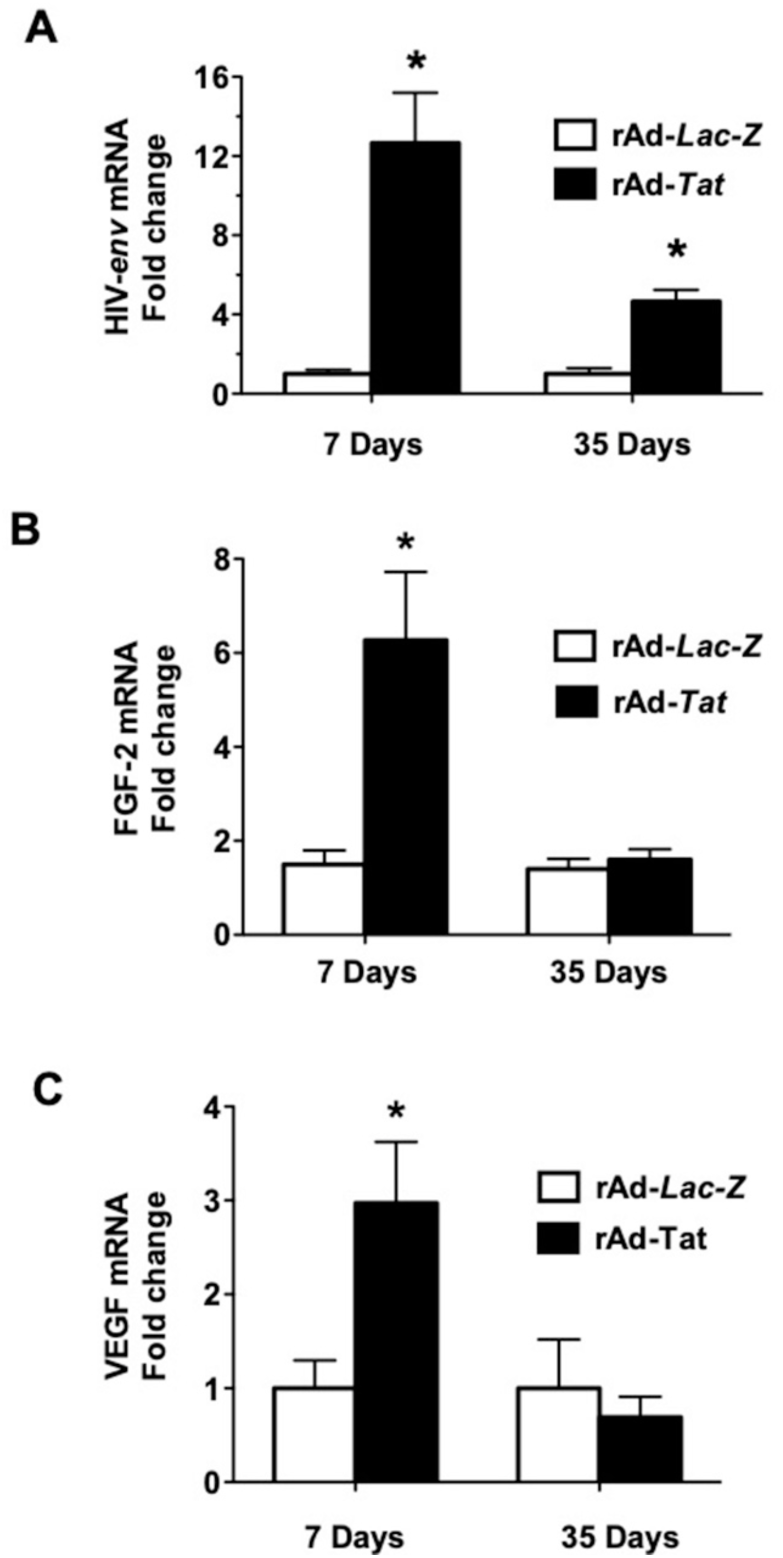
rAd-Tat increased the expression of HIV-envelope (env), Fibroblast Growth Factor-2 (FGF-2), and Vascular Endothelial Growth Factor (VEGF) mRNA. RNA was extracted from the kidney of 7 and 35 days old HIV-Tg_26_ mice infected with rAd-*Lac-*Z or rAd-*Tat* vectors (n = 4-6 mice per group). Real time qRT-PCR analysis of HIV-*env*, FGF-2, and VEGF was done as described in the methods section. * Unpaired *t* test analysis between the two groups at 7 and 35 days of life showed p values < 0.05.

### Tat-induced HIVAN in HIV-Tg_26_ mice

Seven days old HIV-Tg_26_ mice injected with rAd-Tat developed proteinuria and renal histological injury in association with an up-regulated expression of HIV-*env* (Figure 3A-C). In contrast, wild type mice injected with HIV-Tat or *Lac-Z* vectors did not develop significant renal histological lesions or proteinuria by the end of the study period (Figures 4A-B). By 35 days of life, heterozygous HIV-Tg_26_ mice injected with rAd-Tat vectors developed significant proteinuria and HIVAN-like renal histological lesions (Figure 4A-B). Furthermore, the BUN levels of these mice were elevated, when compared to HIV-Tg_26_ mice injected with rAd-Lac-Z vectors (35 ± 1 mg/dl* vs. 24.5 ± 2.6 mean± SEM; *p<0.05). Overall, these findings suggest that Tat plays an important role precipitating HIVAN in HIV-Tg_26_ mice.

**Figure 3.**
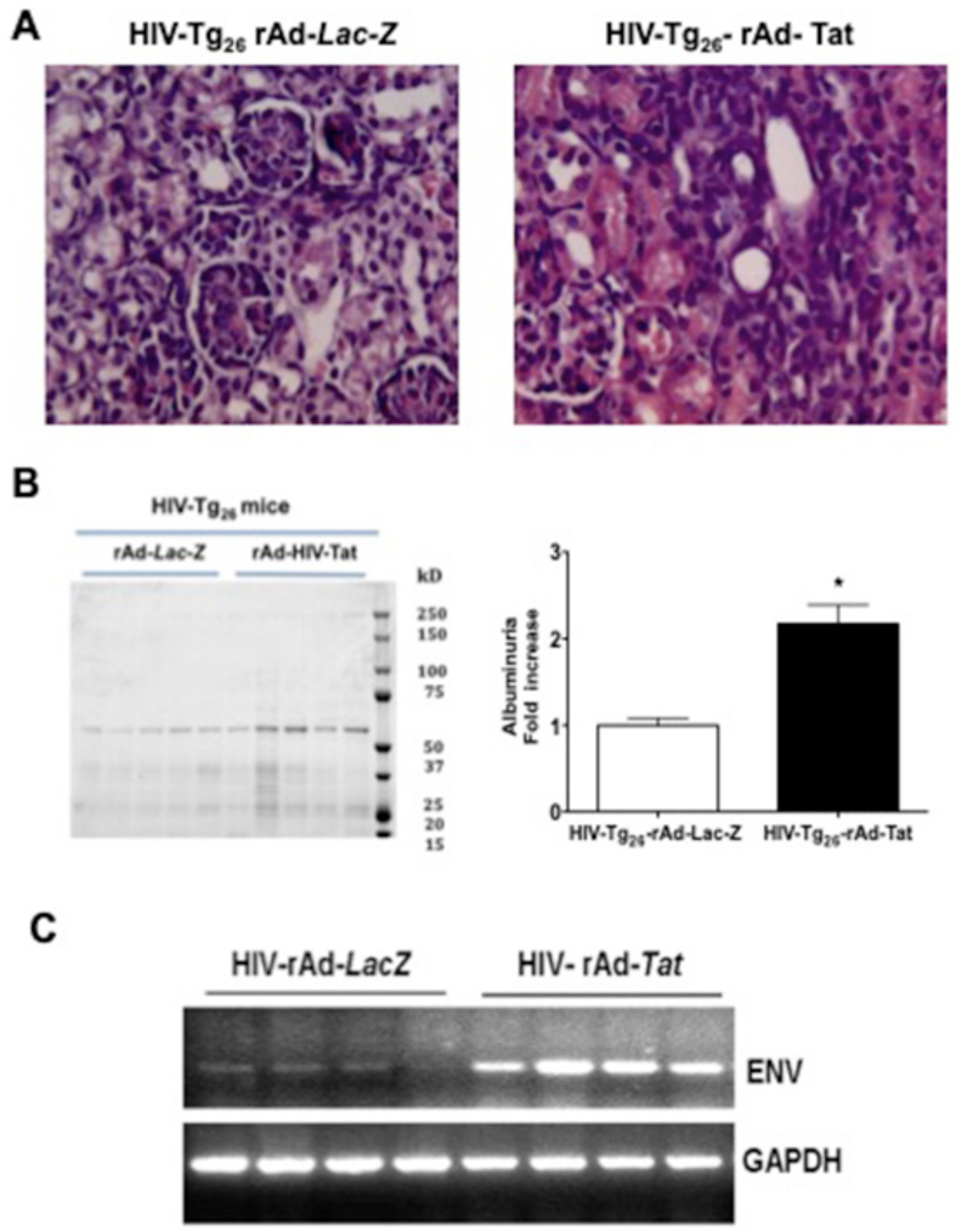
rAd-Tat induced the expression of the HIV-envelope (env) gene in the kidney of 7 days old HIV-Tg_26_ mice, in association with the development of renal histological lesions and albuminuria. **A**. Panel A shows representative renal sections harvested from 7 days old HIV-Tg_26_ mice injected either with rAd-*LacZ* or rAd-*Tat* vectors and stained with hematoxylin & eosin. Original magnification x 250. **B**. Panel B shows a Coomassie blue stained SDS-PAGE gel loaded with urine samples (5 microliters) collected from 7 days old HIV-Tg_26_ mice injected either with rAd-*LacZ* or rAd-*Tat* vectors (n = 5 per group). Quantitation of albuminuria was assessed by densitometric analysis of the Coomassie blue stained albumin bands. Results were expressed in arbitrary optical density (OD) units as a ratio of urinary creatinine and compared to control baseline values. * Unpaired t-test, p < 0.01 (n = 5 mice per group). **C.** Panel C shows the expression of HIV-*envelope* (*env*) and GAPDH mRNA by RT-PCR in the kidney of 7 days old HIV-Tg_26_ mice infected with either rAd-*LacZ* or rAd-*Tat* vectors (n = 4 mice per group).

**Figure 4.**
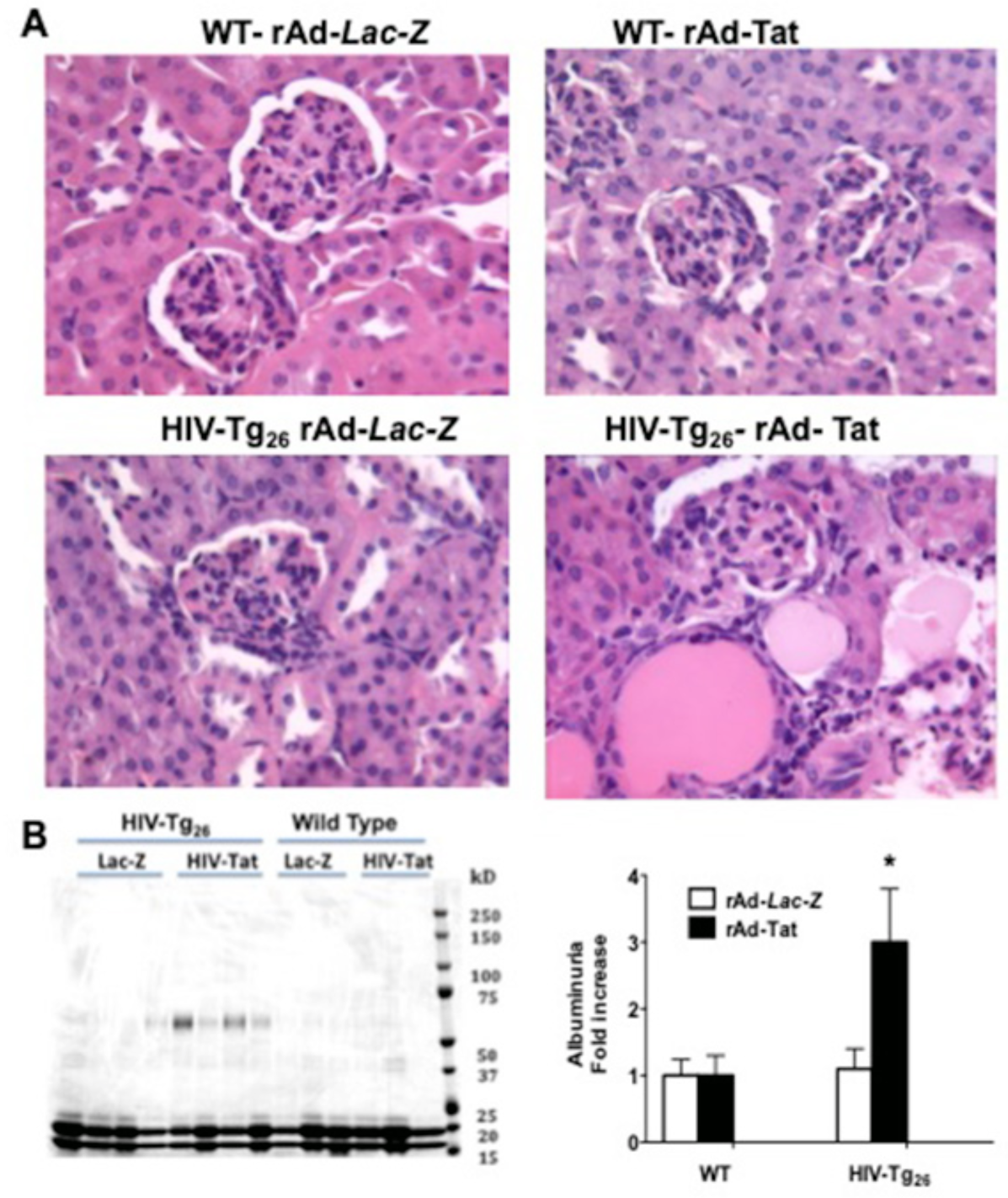
rAd-Tat induced HIV albuminuria AND HIVAN in 35 days old HIV-Tg_26_ mice. **A**. Panel A shows representative renal sections harvested from 35 days old wild type (WT) and HIV-Tg_26_ mice infected with rAd-*LacZ* or rAd-*Tat* vectors and stained with hematoxylin & eosin. Originalmagnification x 250. **B**. Panel B shows a Coomassie blue stained SDS-PAGE gel loaded with urine samples (5 microliters) collected from 35 days old wild type and HIV-Tg_26_ mice infected with either rAd-*LacZ* or rAd-*Tat* vectors (n = 4 - 3 per group). Quantitation of albuminuria was assessed by densitometric analysis of the Coomassie blue stained albumin bands. Results were expressed in arbitrary optical density (OD) units as a ratio of urinary creatinine as described in the methods section. * Unpaired t-test, p < 0.01 (n = 4 mice per group).

### Renal proliferative changes in young HIV-Tg_26_ mice

As shown in Figures 5 – 8, HIV-Tg_26_ mice injected with rAd-Tat develop significant renal epithelial proliferative changes. Briefly, immunohistochemistry and Western blots studies revealed that the expression levels of PCNA, Ki-67, and MAPK were elevated in 7 days old HIV-Tg_26_ mice injected with rAd-Tat, when compared to mice injected with rAd-*Lac-Z* control vectors (Figures 5–6). In contrast, no changes were detected in the kidney of 7 or 35 days old wild type mice injected with rAd-Tat or *Lac-Z* vectors (Supplemental Figures 1–2). Taken together, these findings suggest that HIV-Tat interacts with other HIV-1 genes and/or cytokines to induce the proliferation of renal epithelial cells [15, 16, 18]. Alternatively, using an in situ apoptosis detection kit and Western blots to detect caspase-3 activation, we found a reduced number of renal epithelial cells undergoing apoptosis in 7 days old HIV-Tg_26_ mice injected with rAd-Tat, relative to those infected with the control rAd-*Lac-Z* vectors (Figures 5 – 6). The renal epithelial proliferative changes and MAPK pathway activation were also seen in 35 days old HIV HIV-Tg_26_ mice injected with rAd-Tat (Figures7–8), but no differences in apoptosis or caspase-3 activation were noted at this latter stage (Figure 7).

**Figure 5.**
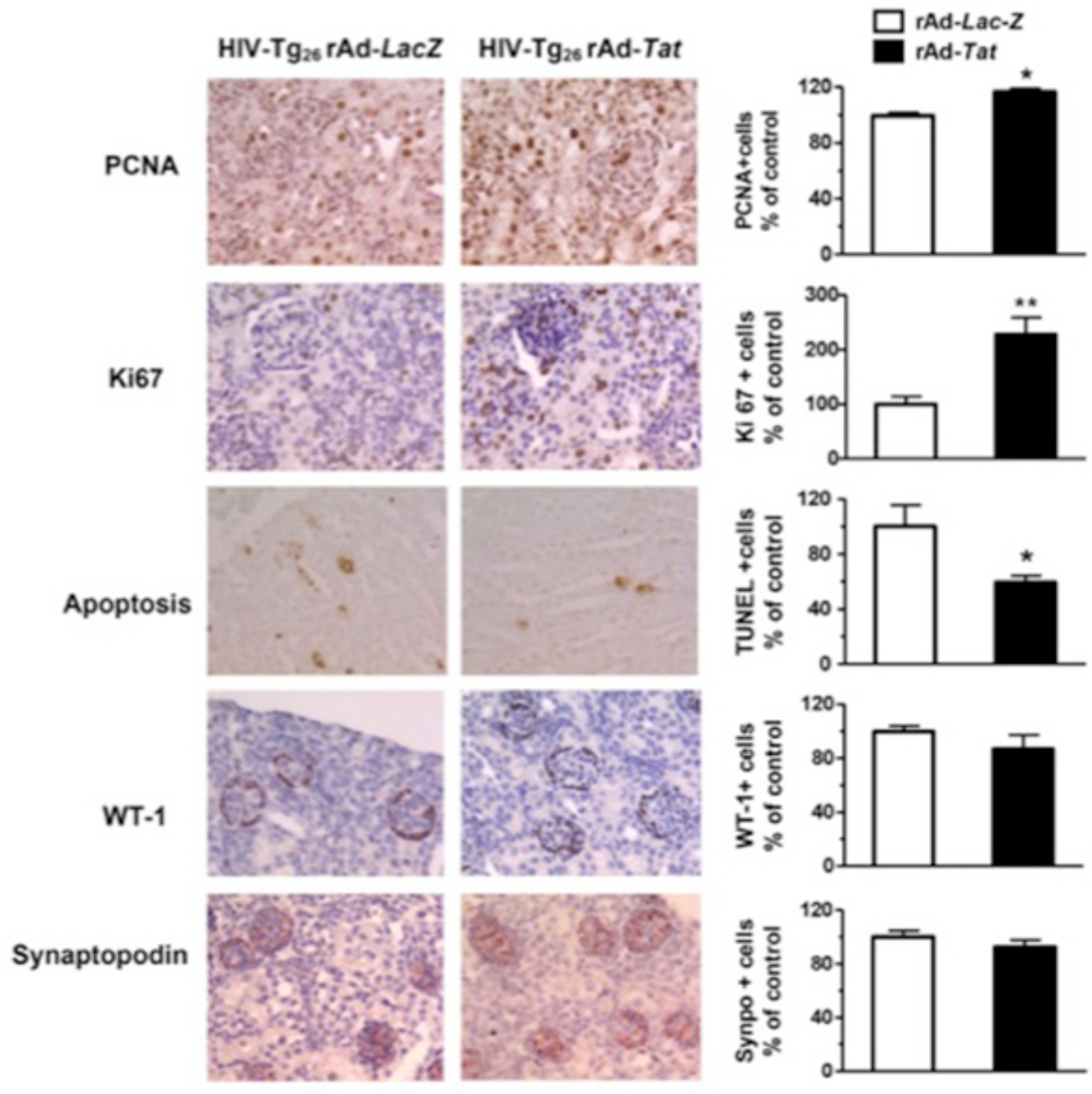
rAd-Tat induced proliferative and anti-apoptotic changes in the kidney of 7 days old HIV-Tg_26_ mice. The panels shows representative immunohistochemistry staining for the proliferating cell nuclear antigen (PCNA) and the Ki-67 antigen (both brown color), apoptosis (brown color), WT-1 antigen (red color), and synaptopodin (red color) in renal sections harvested from 7 days old HIV-Tg_26_ mice infected with either rAd-*Tat* or rAd-*Lac-Z* vectors. The graphs represent percentage changes in positive cells (mean ± SEM) relative to the controls (n = 4 - 5 per group; unpaired t-test * p<0.05 or ** p<0.01 when compared to the HIV-Tg_26_ mice infected with rAd-*Lac-Z* control vectors). Original magnification: X250.

**Figure 6.**
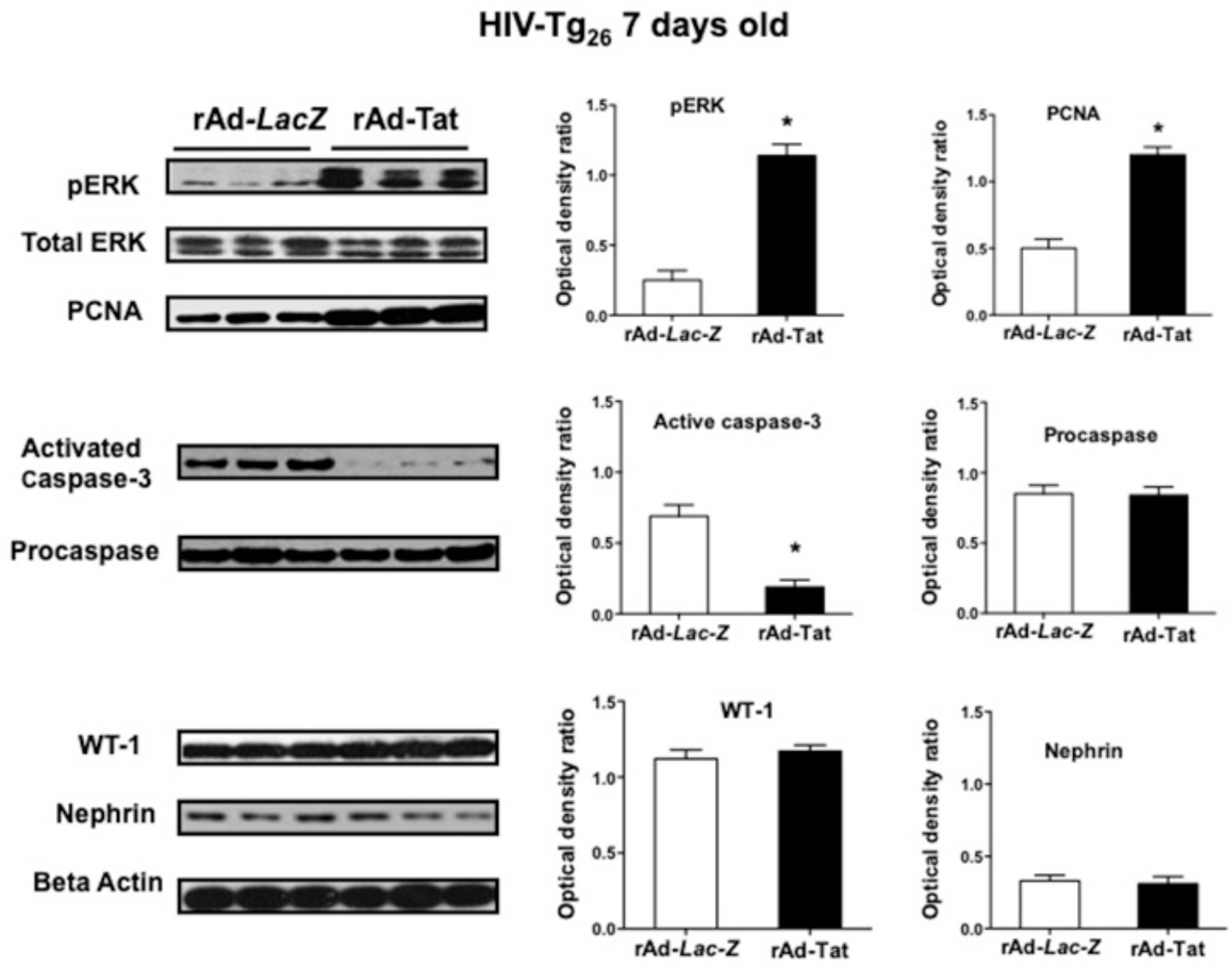
rAd-Tat induced proliferative and anti-apoptotic changes in the kidney of 7 days old HIV-Tg_26_ mice. The panels show representative results of the Western blot analysis for phospho-p44/42 MAPK (p-ERK), proliferating cell nuclear antigen (PCNA), activated caspase-3, procaspase, Wilms tumor 1 (WT1) and nephrin done with kidney homogenates derived from 7 days old HIV-Tg_26_ mice infected with rAd-*Tat* or rAd-*LacZ* vectors (n = 5 mice per group). Comparisons between groups were done by unpaired t-test analysis. The expression of PCNA, WT-1 and nephrin was quantified as a ratio of beta actin. The graphs show the results of the densitometry analysis and quantification of the results in optical density units (mean ± SEM), as described in the methods sections. * p < 0.05 compared to HIV-Tg_26_ mice infected with the rAd-*Lac-Z* control vectors.

**Figure 7.**
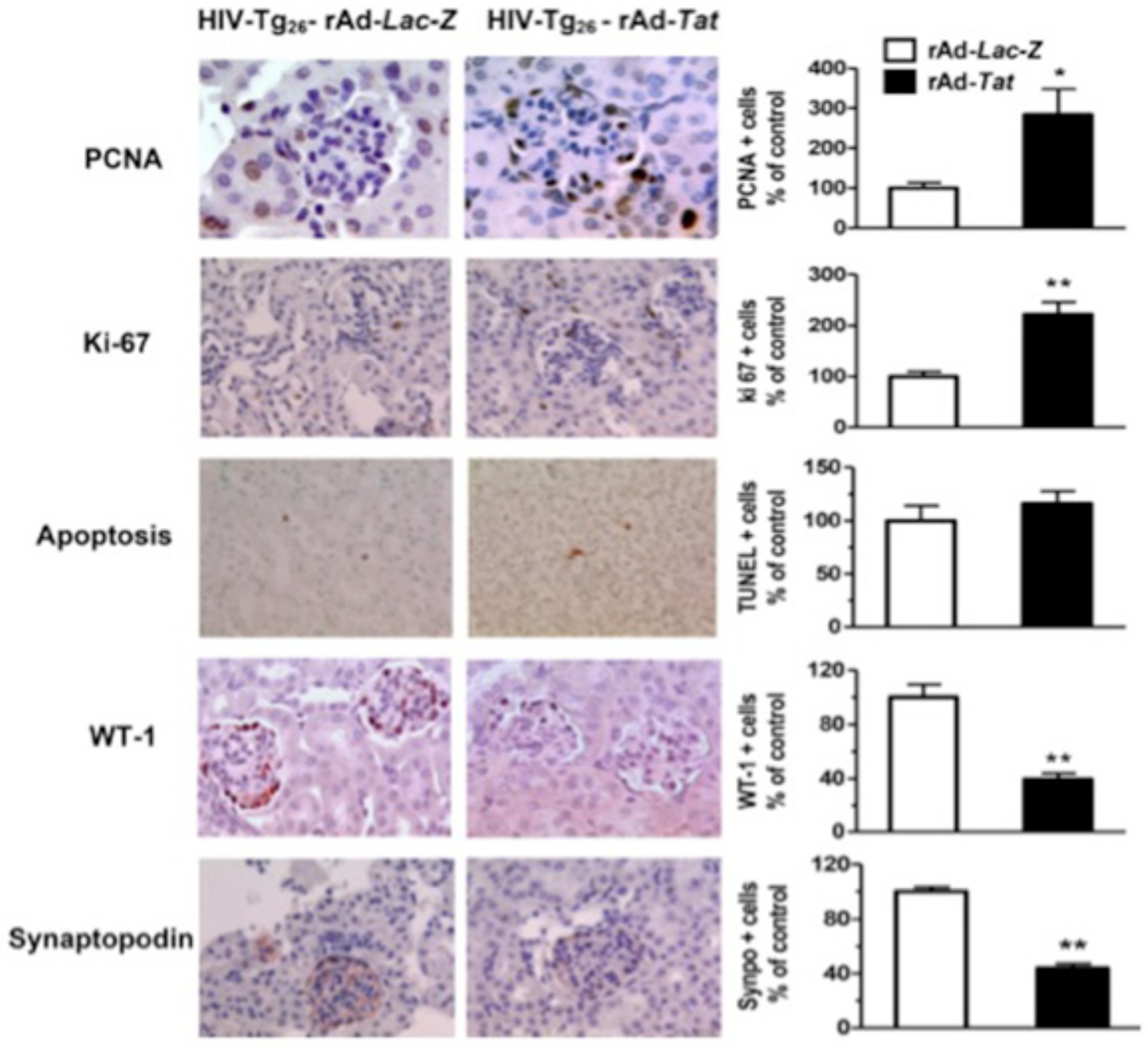
rAd-Tat induced proliferative and de-differentiation changes in podocytes of 35 days old HIV-Tg_26_ mice. The panels shows representative immunohistochemistry staining for the proliferating cell nuclear antigen (PCNA) and the Ki-67 antigen (both brown color), apoptosis (brown color), WT-1 antigen (red color), and synaptopodin (red color) in renal sections harvested from 35 days old HIV-Tg_26_ mice infected with either rAd-*Tat* or rAd-*Lac-Z* vectors. The graphs represent percentage changes in positive cells per field (mean ± SEM) relative to the controls (n = 4 - 5 per group; unpaired t-test * p<0.05 or ** p<0.01 when compared to the HIV-Tg_26_ mice infected with rAd-*Lac-Z* control vectors). Original magnification: X250.

**Figure 8.**
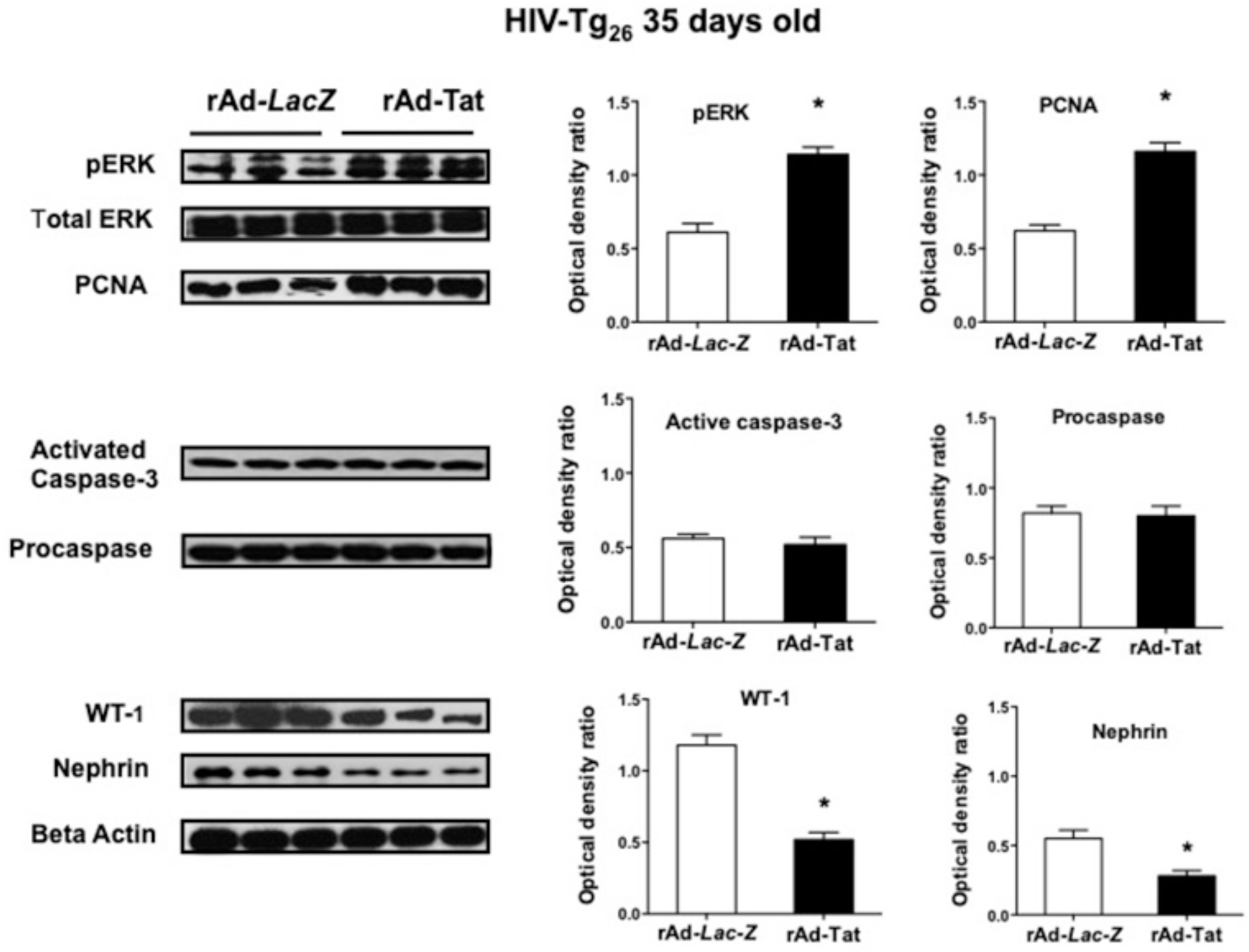
rAd-Tat induced proliferative and de-differentiation changes in podocytes of 35 days days old HIV-Tg_26_ mice. The panels show representative results of the Western blot analysis for phospho-p44/42 MAPK (p-ERK), proliferating cell nuclear antigen (PCNA), activated caspase-3, procaspase, Wilms tumor 1 (WT1) and nephrin done with kidney homogenates derived from 35 days old HIV-Tg_26_ mice infected with rAd-*Tat* or rAd-*LacZ* vectors (n = 5 mice per group). The expression of PCNA, WT-1 and nephrin was quantified using arbitrary optical density units expressed as a ratio of beta actin. The graphs show the results of the densitometry analysis and quantification of the results in optical density units (mean ± SEM), as described in the methods sections. Comparisons between groups was done by unpaired t-test analysis; *p < 0.05, compared to HIV-Tg_26_ mice infected with the rAd-*Lac-Z* control vectors.

### De-differentiation of podocyte in young HIV-Tg_26_ mice

Immunohistochemistry and Western blot studies done in kidney sections derived from 7 days old HIV-Tg_26_ mice infected with rAd-Tat and *Lac-Z* vectors showed no significant differences in the expression levels of the podocyte specific proteins nephrin, WT-1, and synaptopodin (Figure 5–6). However, the protein expression levels of nephrin, WT-1, and synaptopodin were significantly reduced in 35 days old HIV-Tg_26_ mice injected with rAd-Tat vectors (Figures 7–8). In summary, by 35 days of life, almost all HIV-Tg_26_ mice infected with rAd-HIV-Tat vectors develop clinical and renal histological features consistent with HIVAN [1, 15, 19].

## Discussion

In this study we describe a new inducible mouse model system of childhood HIVAN. This model mimics the physiological process by which HIV-1 transcription is activated in humans, and reproduces the full HIVAN phenotype in young HIV-Tg_26_ mice. Our findings underscore the critical role that HIV-Tat plays in the pathogenesis of HIVAN by inducing the renal expression of HIV-1 genes in synergy with heparin-binding growth factors and by increasing the de-differentiation and proliferation of renal epithelial cells.

To develop the mouse model of childhood HIVAN we took advantage of the HIV-Tg_26_ mouse line [7, 15, 19]. These mice carry a 7.4-kb HIV-1 construct lacking 3-kb sequence overlapping the *gag/pol* region of the provirus pNL4-3 [7, 15, 19] and express HIV-1 transcripts in many tissues, including kidney glomerular and tubular epithelial cells. Homozygous HIV-Tg_26_ mice are born sick and usually died with multiple systemic lesions during the first days or weeks of life [7, 19]. In contrast, heterozygous mice can be followed until they reach adulthood, and have been used by several investigators to explore the pathogenesis of HIVAN [7, 15, 16, 19]. Because the majority of heterozygous HIV-Tg_26_ mice develop HIVAN at different time points after they reach adulthood, currently we do not have a reliable mouse model system to study the pathogenesis of childhood HIVAN. Therefore, we carried out this study to test the hypothesis that the induction of HIV-genes in the kidney of newborn mice precipitates HIVAN during the first month of life. To accomplish this goal, we used an adenovirus gene transferring technique developed in our laboratory, which is based on the principle that the retention of adenoviral vectors in the circulation improves the transduction of renal glomerular cells in rodents [20, 21] In previous studies, we showed that newborn mice have delayed clearance of rAd vectors from the circulation, and therefore, more efficient transduction of glomerular cells after a systemic injection of adenoviruses via the retro-orbital plexus [14]. Following this approach, we expressed the coding sequences of a Tat gene derived from a child with HIVAN (Tat-HIVAN) in the kidney of newborn HIV-Tg_26_ mice, and precipitated the development of HIVAN during the first month of life. Our findings support the results of previous studies showing that HIV-1 genes expressed in the kidney play a critical role in this process, although we do not yet understand the exact mechanisms involved. Further studies are warranted to explore this issue.

The HIV-Tat protein is a powerful transcriptional factor encoded by two exons. The first exon encodes the HIV activation and basic binding domains, which are required for HIV-transcription and nuclear localization of Tat [22]. The second exon encodes the RGD motif (C-terminal amino acids 73–86), which enhances the angiogenic activity of Tat acting through cytokines and integrin receptors [23]. Tat plays an essential role in HIV-replication by recruiting a cellular human protein called cyclin T1, which efficiently increases the transcription of the HIV-LTR via NF-κB [24]. However, cloning and characterization of the murine CycT1 protein revealed that mouse cyclin 1 lacks a critical cysteine residue that is needed to form a complex with Tat and induce its full transcriptional activity [25, 26]. For this reason, Tat has limited direct transcriptional activity in mice, but it can induce the expression of TNF-α[27] and other cytokines that increase the transcription of HIV-1 via NF-κB dependent mechanisms [28]. Our study showed that the activation and basic binding domains of Tat are sufficient to induce the renal expression of HIV-genes and precipitate HIVAN in young mice. In contrast, we found Tat’s RGD motif is not essential in this process.

In addition to being a powerful activator of HIV-1 transcription, Tat is released into the circulation by infected cells and can be taken up by uninfected cells [13, 18]. In this manner, Tat mimics the action of several cytokines involved in the pathogenesis of AIDS, including SDF-1α, RANTES, and MIF1-β [13, 18, 29]. Furthermore, acting in synergy with FGF-2, Tat can induce the de-differentiation and proliferation of cultured human podocytes [30–32]. For these reasons, we explored the effects of Tat in wild type mice, but were unable to detect significant renal lesions in these mice. Our findings suggest that Tat alone cannot induce renal disease in wild type mice. However, we should mention that Tat-HIVAN has an incomplete RGD sequence, and its ability to interact with cytokines and integrin receptors *in vivo* might be impaired [12, 13]. Thus, further studies are needed to determine whether Tat proteins containing RDG sequences can cause kidney damage *per se* in wild type mice. We speculate that an additional mechanism through which Tat could precipitate HIVAN in HIV-Tg_26_ mice is by increasing the production and/or activity of TNF-α [27], since high levels of TNF-α are detected in the circulation of HIV-Tg_26_ mice [33], and TNF-α worsens the outcome of HIVAN in adult mice [34]. Alternatively, Tat could act in synergy with FGF-2 and VEGF-A [15, 16, 28, 31], considering that both heparin binding growth factors were up-regulated by Tat in 7 day old HIV-Tg_26_ mice, and have been linked to the pathogenesis of HIVAN in children and HIV-Tg_26_ mice [15, 16, 35]. Finally, the Tat-induced expression of the HIV-Tg_26_ transgene should increase the kidney expression levels of *nef* and *vpr*, and both HIV-genes appear to play a critical role in the pathogenesis of HIVAN [10, 11].

Overall, our mouse model reproduces all the renal histological features characteristic of childhood HIVAN [1, 36]. Interestingly, the expression levels of the podocyte specific proteins, nephrin, WT-1, and synaptopodin, did not change in correlation with the induction of HIV-1 genes during the first week of life. These findings suggest that the podocyte de-differentiation changes characteristic of HIVAN, might be a secondary event associated with the regeneration of these cells. It is possible that podocytes that express high levels of HIV-1 genes died, and were replaced by parietal epithelial or renal progenitor cells [37, 38], which do not express podocytes markers, and are sensitive to the growth promoting effects of several growth factors [39]. In addition, we noted a reduced number of renal epithelial cells undergoing apoptosis in 7 days old HIV-Tg_26_ mice infected with rAd-Tat vectors, when compared to the controls. It is tempting to speculate that Tat, in combination with bFGF-2 and VEGF-A, may have an anti-apoptotic effect [40], since both heparin binding growth factors were up-regulated at this time point. However, more studies are needed to test this hypothesis. Finally, we found that Tat induced direct renal epithelial proliferative changes in 7 and 35 days old HIV-Tg_26_ mice. These changes appear to be driven by the MAPK signaling pathway, which can be activated directly by HIV-Nef [41–44], as well as FGF-2 [15] or VEGF-A [16]. In summary, by the end of the first month of life, all HIV-Tg_26_ mice infected with rAd-Tat vectors develop proteinuria and renal histological lesions consistent with HIVAN.

In humans, the risk variants of the Apolipoprotein-1 (APOL1) increase the lifetime risk of untreated HIV+ people to develop HIVAN by ~ 50% [45–47]. Therefore, one limitation of our animal model is that HIV-Tg_26_ mice do not express the APOL-1 gene. This limitation could be overcome by generating dual transgenic HIV-Tg_26_ / APOL1 mice [48], and infecting newborn mice with rAd-Tat vectors. In addition, a significant number of Black children living with HIV develop HIVAN independently of the APOL1 risk variants [49, 50], and previous studies suggest that the APOL1 risk variants may play more relevant role in adults, when compared to young children [50, 51]. Thus, young kidneys might be more sensitive to the cytotoxic effects of HIV-1 genes, TNF-α, and heparin binding growth factors, and less dependent on the APOL1 risk variants to develop HIVAN. Alternatively, it is possible that other unknown genetic factors may play an additional role precipitating HIVAN in Black children, as reported in HIV-Tg_26_ mice [52]. Taken together, these studies show that a strong genetic influence modulates the outcome of HIVAN both in mice and humans, and more work needs to be done to define these factors in children.

In conclusion, we have developed an inducible mouse model system of childhood HIVAN that reproduces the full HIVAN phenotype during their first month of life. In addition, we showed that Tat plays a relevant role in this process by inducing the renal expression of HIV-1 genes, FGF-2, and VEGF-A, leading to the activation of the MAPK pathway. Hopefully, this animal model will facilitate the discovery of new therapeutic targets to prevent the progression of HIVAN in children and adolescents.

## Materials and Methods

### Experimental design

This study was approved by the Children’s Research Institute Animal Care and Use Committee. Heterozygous newborn HIV-Tg_26_ FVB/N mice [7, 19] and their respective wild type (WT) littermates were injected through the retro-orbital plexus with 2 × 10^9^ pfu/mouse of recombinant adenoviral (rAd) vectors carrying either HIV-Tat derived from a child with HIVAN (rAd-*Tat* vector) or the *E. coli LacZ* gene (rAd-*Lac-Z*). To express the HIV-Tat rAd vector in the kidney of newborn mice, we used a gene transferring technique developed in our laboratory [14]. Wild type and HIV-Tg_26_ mice expressing the pro-viral DNA construct d1443 [15, 19], were divided in groups (n = 8 mice each) and sacrificed at 7 days (peak of rAd-Tat expression) and 35 days (renal clearance of the viral vectors) after the adenoviral injections. All mice had free access to water and standard food and were treated in accordance with the National Institutes of Health (NIH) guidelines for care and use of research animals.

### Adenoviral vectors

The generation of the rAd-Tat vector derived from a child with HIVAN was described in detail before [31]. Briefly, a cDNA fragment encoding the full-length Tat protein was cloned into the pCXN2-FLAG vector and used to generate E1-deleted recombinant adenoviruses carrying Tat-HIVAN-FLAG [31]. The protein sequence of the Tat-HIVAN gene was aligned and compared to other Tat genes using the Clustal Omega multiple sequence alignment program (http://www.ebi.ac.uk/Tools/msa/clustalo/). Both Tat-FLAG and *Lac-Z* control adenoviruses were amplified, purified, desalted, and titrated as previously described [14, 53, 54]. The particle-to-plaque-forming unit (pfu) ratio of the virus stock used in these experiments was 100.

### Blood, urine and kidney sample collection

Mice were sacrificed 7 and 35 days after the rAd injections. Urine, blood and kidney samples were harvested and kept frozen at −80°C. Blood urea nitrogen was assessed using the Quantichrom Urea Assay Kit from BioAssay Systems (Catalog No. DIUR-500) as described before [55]. The urinary creatinine levels were measured using colorimetric assay from R&D Systems (Catalog No. KGE005). Urinary protein was measured using the Bayer Multistix 10 SG reagent strips for urinalysis. In addition, 5 microliters of urine were run on 4-12% sodium dodecyl sulfate polyacrylamide gel electrophoresis (SDS-PAGE) and stained with Coomassie Blue Stain Solution (Bio-Rad) to detect proteinuria. The protein band corresponding to albumin was quantified by densitometry analysis using Adobe Photoshop 6.0 (Adobe Systems, San Jose, CA). Results were expressed in arbitrary optical density units adjusted to the urinary creatinine values as described before [32].

### RT-PCR analysis

Total kidney RNA was isolated using Trizol (Invitrogen, Catalog No.15596-026), and treated with deoxyribonuclease I (Dnase I) following Invitrogen’s protocol for RT-PCR studies. cDNA was generated from 3 µg RNA using SuperScript III First-Strand Synthesis System for RT-PCR (Invitrogen, Catalog No. 18080-051). Tat mRNA expression was assessed by RT-PCR using the following primers: forward 5′-ATG GAG CCA GTA GAT CCT AGAC-3′ and reverse 5′-CTA ATC GAA TCG ATC TGT CTC TGC-3′. To determine the relative expression of HIV-1 *envelope (env)* we used the following primers: forward primer 5’-TGT GTA AAA TTA ACC CCA CTC TG-3’, and the reverse primer 5’-ACA ACT TAT CAA CCT ATA GCT GGT-3’. As a control we amplified the mouse housekeeping gene glyceraldehydes-3-phosphate dehydrogenase (*Gapdh*) using the forward primer 5’-CTT ACT CCT TGG AGG CCA TGT-3’, and the reverse primer 5’-GCC AAG GTC ATC CAT GAC AAC-3’). During the amplification process, samples were kept at 94°C for 4 min, followed by 35 cycles at 94°C for 30s, 55°C for 30s, 72°C for 1 min, and a final extension of 8 min. For each HIV-envelope and GAPDH PCR amplification reaction, we used 5µl and 2µl cDNA respectively. The densitometry analysis was conducted using Adobe Photoshop 6.0 as described before [31, 56].

### Real-time RT-PCR analysis

Real-time RT-PCR studies were performed on cDNA samples using the Platinum qPCR SuperMix-UDG kit (Catalog No.11730-017, Invitrogen). The HIV-envelope assay was designed to amplify a 95-bp amplicon from HIV-1 NL4-3 (GenBank accession no. AF324493). Forward primer 5’-CCT TTG AGC CAA TTC CCA TAC ATT-3, reverse primer 5’-gacgttTGG TCC TGT TCC ATT GAA CGT C-3’ with FAM-labeled LUX. The mouse nephrin assay was designed to amplify a 79-bp amplicon (GenBank accession no. NM_019459.2). Forward primer 5’-GTC GGA GGA GGA TCG AAT CAG-3’, reverse primer 5’-cgggGTG GAG CTT CTT GTG TCC CG-3’ with FAM-labeled LUX. The mouse fibroblast growth factor 2 (*Fgf2*) assay was designed to amplify a 70bp amplicon (GenBank accession no. NM_008006), forward primer 5’-CCG GTC ACG GAA ATA CTC CAG-3’ forward primer, reverse primer 5’-cgaactCCG AGT TTA TAC TGC CCA GTT CG-3’ with FAM-labeled LUX (Cat. no19450335, Invitrogen). The mouse Vascular Endothelial Growth Factor assay, VEGF_164_ isoform, was designed to amplify a 101-bp amplicon (GenBank accession no. M95200.1), forward primer 5’-cggcCTA CCA GCG AAG CTA CTG CCG-3’ with FAM-labeled LUX, reverse primer 5’-CAC ACA GGA CGG CTT GAA GAT G-3’. The mouse glyceraldehydes-3-phosphate dehydrogenase (*Gapdh*) housekeeping gene control assay, was designed to amplify a 93 bp amplicon from (GenBank accession no. NM_008084.1). Forward primer 5’-gacatacAGG CCG GTG CTG AGT ATG T-3’ with JOE-labeled LUX, reverse primer 5’-TTT GGC TCC ACC CTT CAA GT-3’. The real-time PCR amplification protocol was as follows: 50°C, 2 min hold (UDG treatment), 95°C, 2 min and 40 cycles of 95°C, 15s, 58°C, 30s, and 72°C, 30s using a 7900 Fast Real-Time PCR System (AB Applied Biosystems). Data were normalized to *Gapdh* and presented as fold increase compared to the rAd-*Lac-Z* control group.

### Western blot analysis

The kidneys were lysed using RIPA lysis buffer containing protease inhibitors and phosphatase inhibitor cocktail 2 (Sigma-Aldrich), and processed by Western blots as described before [3]. The following primary antibodies were used: phospo-p44/42 mitogen-activated protein kinase (Thr202/Tyr204) p44/42 mitogen-activated protein kinase ERK1/2, both obtained from Cell Signaling Technology, Proliferating cell nuclear antigen (PCNA) (C-20) rabbit polyclonal; β actin (I-19) goat polyclonal, caspase-3 (pro and activate forms) rabbit polyclonal antibodies (Santa Cruz Biotechnology), Wilms’ Tumor 1 (WT-1) mouse monoclonal anti-human antibody, clone 6F-H2 (Dako North America, Inc), Nephrin guinea pig polyclonal antibody (Progen Biotechnik GmbH). All primary antibodies were diluted 1:1000 except WT-1, which was diluted 1:500 dilution and incubated overnight at 4°C. Protein bands were detected using Supersignal West Pico Chemiluminescent Substrate (Thermo Scientific) according to the manufacturer’s instruction. All membranes were exposed to a Kodak film (X-OMAT) and developed using an automated developer. Densitometry analysis of the data expressed as a β actin ratio was conducted using Adobe Photoshop 6.0 as described before

### Immunohistochemistry

Paraffin embedded sections were cut at 5μm, deparaffinized, rehydrated, and stained as previously described [54]. Immunostaining was performed with a commercial streptavidin-biotin-peroxidase complex (Histostain SP kit, Zymed, San Francisco, CA) according to the manufacturer’s instructions as described before [57]. The peroxidase activity was monitored after the addition of substrate using a DAB kit (Vector Laboratories, Catalog No SK-4100) or AEC substrate kit (Catalog No. 002007,Invitrogen, Frederick, MD). Sections were counterstained with Hematoxylin. The PCNA staining kit from Invitrogen was used to detect PCNA. Ki-67 and Wilms’ Tumor 1 (WT-1) staining were assessed using a 1:50 dilution of a monoclonal rat anti-mouse Ki-67 antibody (clone TEC-3) and a mouse monoclonal anti-human WT-1 antibody (clone 6F-H2) respectively, both from Dako North American Inc). Synaptopodin was detected with a ready-to-use a mouse monoclonal antibody (clone G1D4, Batch No. 1372) from Fitzgerald Industries International, INC. Controls included replacing the primary antibody with equivalent concentrations of the corresponding non-specific antibodies and/or omitting the first or second antibodies. Apoptosis was assessed using the Apop Tag *in situ* apoptosis detection kit (Catalog No S7101) from Chemicon International, according to the manufacturer’s instructions.

### Statistical analysis

If not specified otherwise, the data were expressed as mean + SEM. Differences between two groups were compared using an unpaired *t* test. Multiple sets of data were compared by ANOVA with Newman-Keuls *post-hoc* comparisons. Statistical analysis were performed using Graph Pad Prims software (Version 5.00; Graph Pad Software, San Diego, CA. Values of p < 0.05 were considered statistically significant.

## Acknowledgements

We thank Dr. Marina Jerebtsova PhD. for her advice expanding adenoviral vectors and perfoming injections in mice, and Dr. Xuefang Xie PhD. for her contribution performing the sequencing and analysis of the Tat-HIVAN gene.

## Supplemental Figures

**Supplemental Figure 1.**
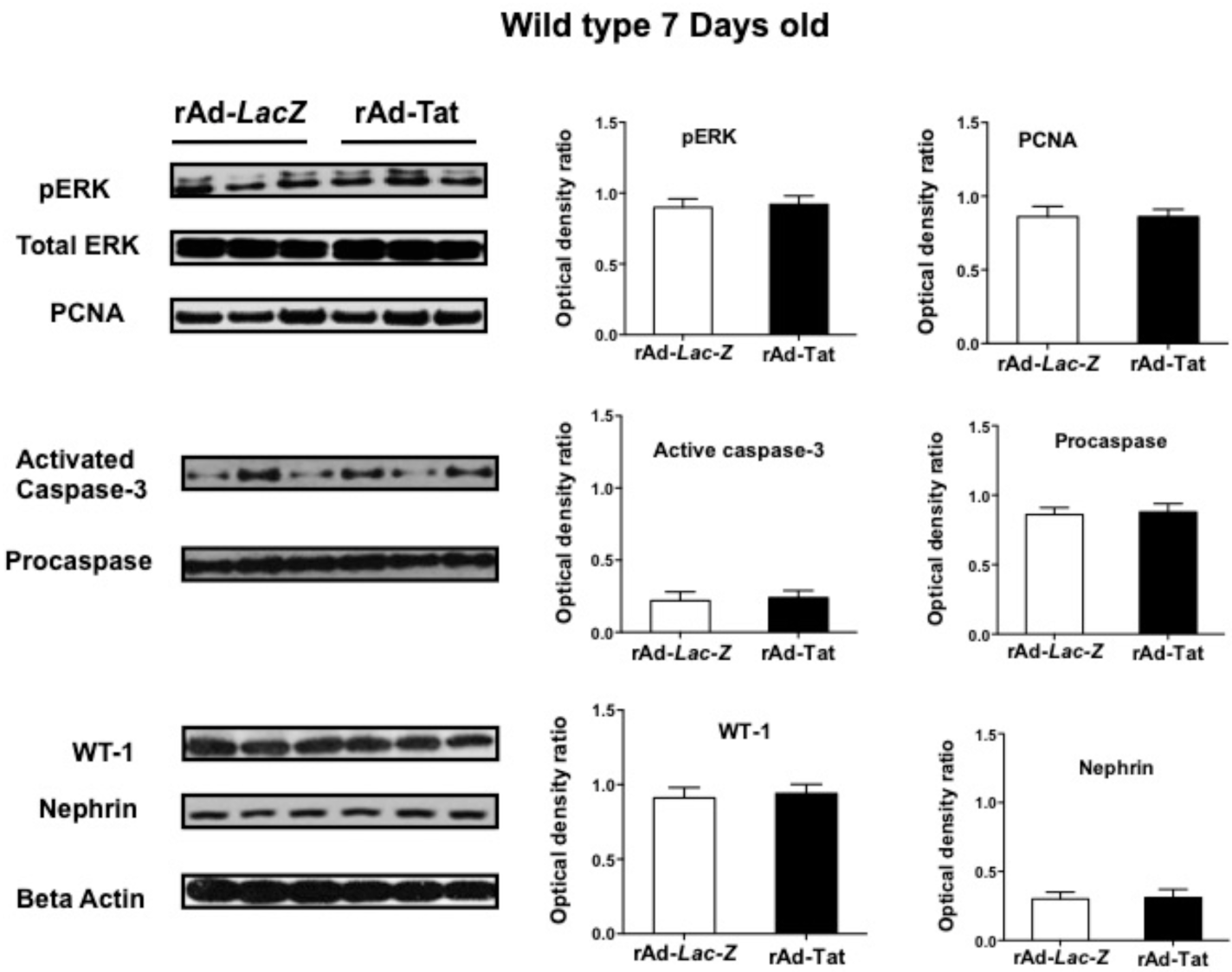
rAd-Tat did not induce significant proliferative or anti-apoptotic changes in the kidney of 7 days old wild type mice. The panels show representative results of the Western blot analysis for phospho-p44/42 MAPK (p-ERK), proliferating cell nuclear antigen (PCNA), activated caspase-3, procaspase, Wilms tumor 1 (WT1) and nephrin done with kidney homogenates derived from 7 days old wild type mice infected with rAd-*Tat* or rAd-*LacZ* vectors (n = 4 mice per group). Comparisons between groups were done by unpaired t-test analysis. The expression of PCNA, WT-1 and nephrin was quantified as a ratio of beta actin. The graphs show the results of the densitometry analysis and quantification of the results in optical density units (mean ± SEM), as described in the methods sections. All changes between groups were not statistically significant (p > 0.05).

**Supplemental Figure 2.**
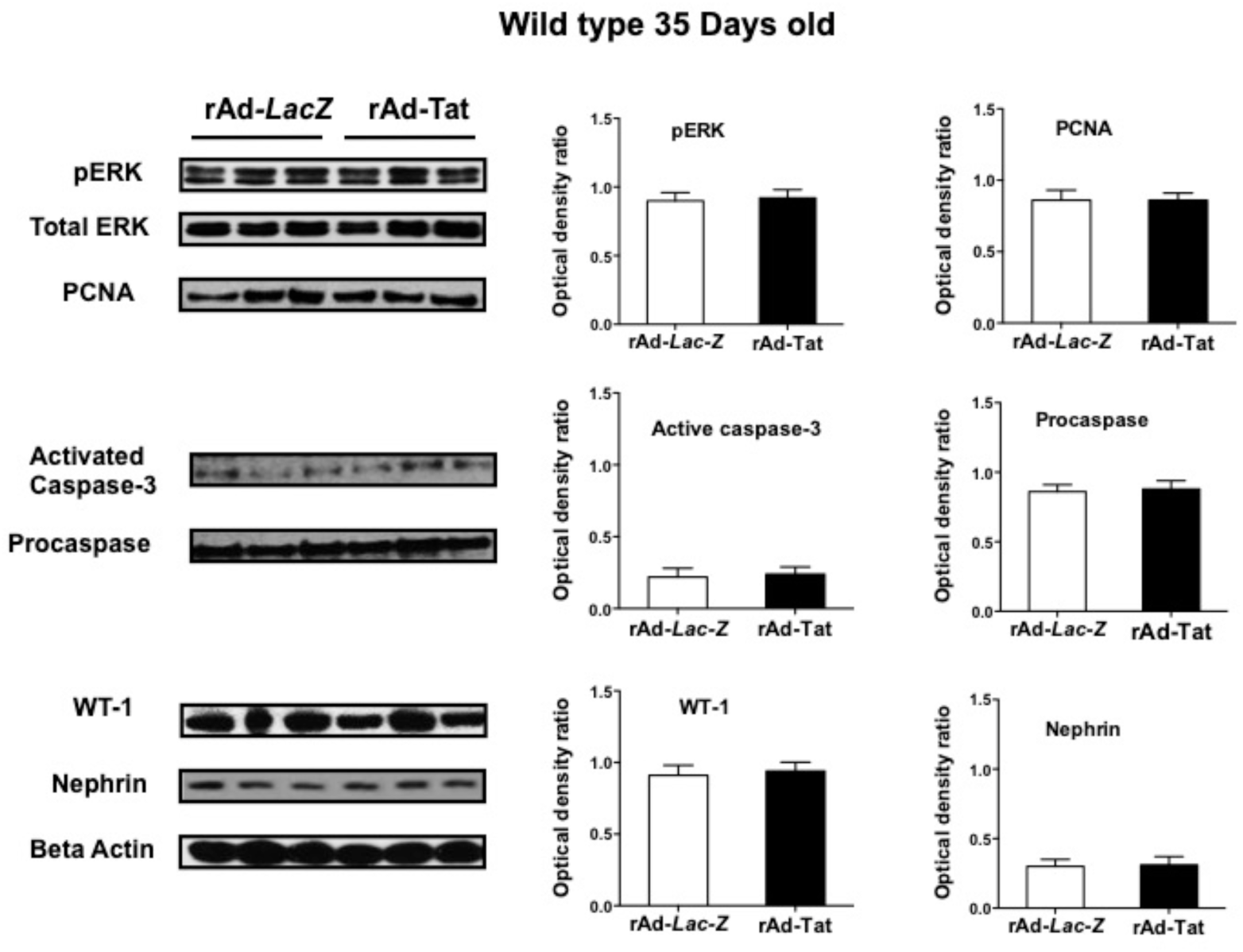
rAd-Tat did not induce significant proliferative or anti-apoptotic changes in the kidney of 35 days wild type mice. The panels show representative results of the Western blot analysis for phospho-p44/42 MAPK (p-ERK), proliferating cell nuclear antigen (PCNA), activated caspase-3, procaspase, Wilms tumor 1 (WT1) and nephrin done with kidney homogenates derived from 35 days old wild type mice infected with rAd-*Tat* or rAd-*LacZ* vectors (n = 4 mice per group). Comparisons between groups were done by unpaired t-test analysis. The expression of PCNA, WT-1 and nephrin was quantified as a ratio of beta actin. The graphs show the results of the densitometry analysis and quantification of the results in optical density units (mean ± SEM), as described in the methods sections. All changes between groups were not statistically significant (p > 0.05).

